## Supplemental legends and figures for "Inner hair cell synapse density influences auditory processing"

**Fig, S1. Gap inhibition is stable across different sessions for all genotypes.** The relationship between gap inhibition vs. gap length does not change between the three timepoints for all genotypes. Mean ± SEM are shown.

**Fig, S2. The degree of gap inhibition correlates with ABR peak I.** The amplitude of ABR peak I versus gap inhibitory level of Ntf3-KD and their littermate controls (**A**) or Nft3-OE and their littermate controls (**B**) show a linear correlation.

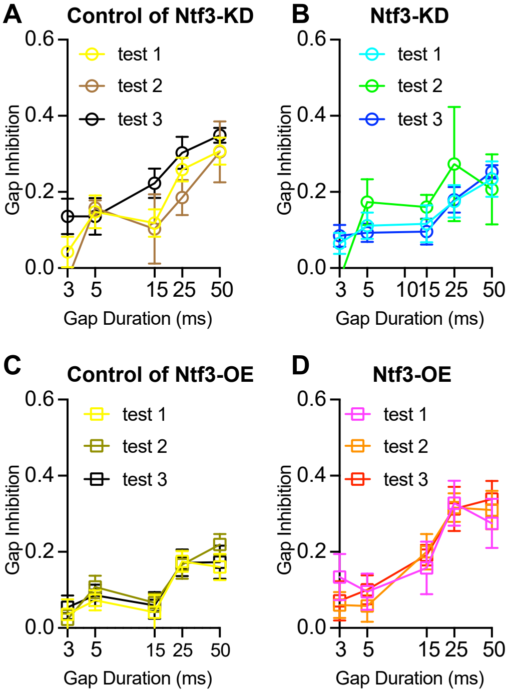
A
Control of Ntf3-KD
B
Ntf3-KD
C
Control of Ntf3-OE
D
Ntf3-OE
test 1
test 2
test 3
Gap Inhibition
Gap Duration (ms)
0.0
0.2
0.4
0.6
3
5
10
15
25
50

Fig. S1

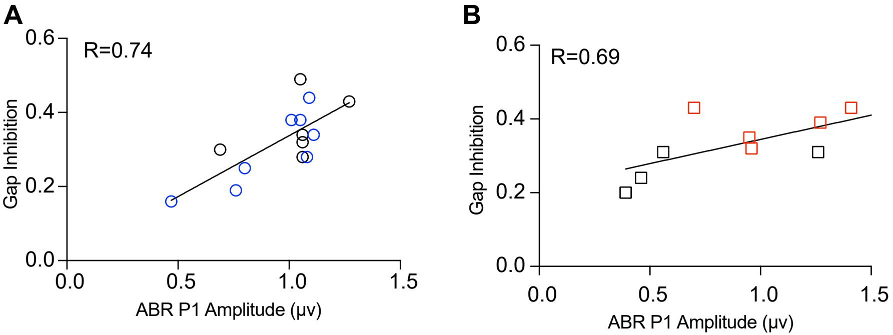
A
R=0.74
Gap Inhibition
0.0
0.2
0.4
0.6
0.0
0.5
1.0
1.5
ABR P1 Amplitude (μv)
B
R=0.69
Gap Inhibition
0.0
0.2
0.4
0.6
0.0
0.5
1.0
1.5
ABR P1 Amplitude (μv)

Fig. S2
